# Complex-centric proteome profiling by SEC-SWATH-MS

**DOI:** 10.1101/323683

**Authors:** Moritz Heusel, Isabell Bludau, George Rosenberger, Robin Hafen, Max Frank, Amir Banaei-Esfahani, Ben C. Collins, Matthias Gstaiger, Ruedi Aebersold

## Abstract

Proteins are major effectors and regulators of biological processes that can elicit multiple functions depending on their interaction with other proteins. The organization of proteins into macromolecular complexes and their quantitative distribution across these complexes is, therefore, of great biological and clinical significance.

In this paper we describe an integrated experimental and computational technique to quantify hundreds of protein complexes in a single operation. The method consists of size exclusion chromatography (SEC) to fractionate native protein complexes, SWATH/DIA mass spectrometry to precisely quantify the proteins in each SEC fraction and the computational framework *CCprofiler* to detect and quantify protein complexes by error-controlled, complex-centric analysis using prior information from generic protein interaction maps.

Our analysis of the HEK293 cell line proteome delineates 462 complexes composed of 2127 protein subunits. The technique identifies novel subcomplexes and assembly intermediates of central regulatory complexes while assessing the quantitative subunit distribution across them. We make the toolset *CCprofiler* freely accessible, and provide a web platform, *SECexplorer*, for custom exploration of the HEK293 proteome modularity.

**Highlights:**

- Introduction of the concept of complex-centric proteome profiling
- Development of *CCprofiler*, a software framework for complex-centric data analysis
- Detection and quantification of subunit distribution of 462 distinct protein complexes containing 2127 proteins from a SEC-SWATH-MS dataset of HEK293 cells, and identification of novel complex variants such as assembly intermediates
- Statistical target-decoy model to estimate accurate false discovery rates for complexes quantified by complex-centric analysis
- *SECexplorer*, an online platform to support custom complex-centric exploration of SEC-SWATH-MS datasets.

## Introduction

Molecular life science research over the last decades has been transformed by technological advances that aim at exploring biological processes as complex systems of interacting molecules. A range of high throughput technologies to analyze genomes, transcriptomes, metabolomes and proteomes now provide accurate molecular inventories of biological samples at high throughput. Yet, the notion of a modular biology^1^ states that for the definition of the functional state of a cell the organization of cellular molecules into functional modules is as important as the composition of the respective “omes”. This notion has been supported by decades of research into the structure and function of specific macromolecular complexes but the task to systematically probe the organization of biomolecules in the cell has remained technologically challenging. Among all macromolecular modules those containing or consisting of proteins are particularly functionally important because they catalyze and control the vast majority of biochemical functions and constantly adapt to and determine the state of the cell.

For high throughput analytical techniques to generate data sets that are quantitative, reproducible and contain low error rates it has frequently been useful to use prior information to guide the acquisition or analysis of the respective data^2^. For mass spectrometry-based proteomics, the concept of peptide-centric analysis^3^ uses reference fragment ion spectra as prior information to detect and quantify proteolytic peptides in complex samples as surrogates for their corresponding proteins. Peptide-centric analyses have been implemented at a moderate level of multiplexing (tens to few hundred proteins) via selected reaction monitoring (SRM^4^) and parallel reaction monitoring (PRM^5^). More recently, massively parallel data independent analysis strategies (DIA) exemplified by SWATH-MS have been developed that reproducibly quantify tens of thousands of peptides from single sample injections into a mass spectrometer^6–8^. In this manuscript we describe and implement the concept of complex centric analysis. It is intended to systematically detect protein complexes in biological samples and to quantify the distribution of proteins across protein complex instances. Complex-centric analysis uses generic protein interaction information as prior information and conceptually extends the principles of peptide-centric analysis to the level of protein complexes.

Complex-centric proteome profiling consists of the robust and proven technique of size exclusion chromatography (SEC) to fractionate native protein complexes, SWATH/DIA mass spectrometry to precisely and reproducibly quantify proteins across SEC fractions and a new computational analysis strategy implemented in *CCprofiler. CCprofiler* carries out fast and automated detection of protein complexes in datasets of quantitative protein maps from consecutive SEC fractions and controls error rates by means of a target-decoy based statistical model. It uses prior information from generic protein interaction maps to detect and quantify protein complexes in the sample. Complex-centric protein profiling is a new implementation of the general concept of protein correlation profiling^9–11^ that distinguishes itself from earlier implementations^12–14^ by the following: i) the use of SWATH-MS for the data generation provides complete protein elution profiles for each detected protein at quantitative accuracy and a wide dynamic range supporting the quantification of even minor components of the proteome, ii) the development of a statistical model in *CCprofiler* that uses a target/decoy model to calculate a FDR for detected complexes, and iii) the use of prior information from generic protein interaction maps to reduce the erroneous assignment of co-eluting proteins to a complex.

A range of generic protein complex compendia have been generated by different approaches that can be used as prior information for complex-centric analysis. They include: i) the CORUM reference database of complexes^15^ generated by curating results from classical biochemical and biophysical analyses of protein complexes. CORUM presently contains 1753 distinct models of human complexes consisting of 2532 proteins; ii) The BioPlex network^16^ and related protein interaction databases, generated by the mass spectrometric identification of proteins co-purifying with affinity tagged “bait” proteins (AP-MS). BioPlex v1.0 describes 23,744 interactions among 7,688 proteins identified as interactors of 2594 bait proteins; iii) the STRING database^17^, an organism-wide protein-protein interaction network generated by the computational integration of multiple lines of evidence for physical and functional associations. STRING (v10) contains 383,626 high confidence interactions (score ≥ 900) among 10,248 human proteins, and iv) protein complex databases generated by correlation profiling of extensive chromatographic co-fractionation of native complexes, followed by DDA mass-spectrometry ^12–14^. In combination, these interaction compendia constitute an extensive, yet incomplete representation of the organization of the (human) proteome into functional complexes and thus provide an essential resource for the implementation of the complex centric analysis strategy that is supported by the computational framework *CCprofiler*.

We benchmark the method, including the *CCprofiler* algorithm, against a manually curated set of protein complexes and evaluate its complex identification performance against a reference method consisting of multidimensional co-fractionation of native extracts and DDA of individual fractions^12^. The results demonstrate high performance of the *CCprofiler* algorithm in relation to manual benchmarking, with observed true positive rates of up to 91 % (high quality signals) at an FDR of 5 %. The data further shows superior performance of the complex-centric approach in recalling protein complexes compared to the reference method, achieved at a significantly reduced experimental effort (81 vs. 1,163 fractions analyzed by LC-MS/MS). We applied the complex-centric proteome profiling strategy to quantify complexes in a native extract from HEK293 cells in exponential growth state. The results indicate that 55 % of the protein mass is present in the form of complexes that distribute across distinct states of complex formation. The data indicated quantitative complex signals for 462 cellular assemblies if prior knowledge from the CORUM, BioPlex and StringDB reference databases was used and the results were cumulatively integrated. The utility of quantifying the distribution of specific proteins across different resolved submodules is exemplified by the identification of previously unknown substructures of cellular effector complexes such as the proteasome. Finally, we describe and provide access to *SECexplorer*, an interactive online platform for customized expert interpretation of quantitative co-fractionation protein profiles generated by SEC-SWATH-MS. We expect that the complex centric analysis method, the SEC-SWATH dataset representing the organization of the proteome of the cycling HEK 293 cell line and the computational tools to explore the data will find wide application in life science research.

## Results

### Principles and main features of complex-centric proteome analysis

We describe an integrated mass spectrometric and computational method to systematically quantify the modular organization of the proteome. The method is schematically illustrated in Figure 1A and consists of five consecutive steps. First, complexes are isolated from a biological sample under mild conditions that retain their native form and fractionated according to their hydrodynamic radius via high-resolution size exclusion chromatography (SEC). Second, collected, consecutive fractions are subjected to bottom-up mass spectrometric analysis using SWATH/DIA mass spectrometry. Collectively, the thus generated 81 SWATH/DIA maps constitute the dataset that will ultimately be explored by complex-centric analysis of protein SEC elution profiles (Step 5). To accurately quantify protein elution along the SEC chromatographic fractions, peptides are identified and quantified from the composite SWATH/DIA data set in step three by peptide-centric analysis^7,18,19^. Specifically, peptide query parameters for tens of thousands of peptides are generated from a reference spectral library and systematically queried across the dataset to quantify each target peptide in each fraction. Fourth, *CCprofiler* is used to infer quantitative protein elution profiles from the peptide elution profiles across SEC fractions (for details on peptide detection and protein inference along the chromatographic fractions see Figure S1). Fifth, the protein SEC elution profiles are explored via complex-centric analysis using *CCprofiler* along with prior protein interaction information, to detect distinct protein modules and to determine the likelihood that each detected module is correctly identified. Specifically, the complex-centric analysis of *CCprofiler* in steps four and five entails (Fig 1B) (i) protein quantification, (ii) target complex query set generation based on prior protein connectivity information, (iii) the generation of corresponding decoy complex query sets used for downstream error estimation, (iv) detection of complex component subunit co-elution signals along SEC fractions, (v) decoy-based generation of a null model and according error estimation and, (vi) compilation of the results into a report detailing unique, chromatographically resolved instances of complexes and the distribution of shared protein subunits across them (For details, see extended experimental procedures).

**Figure 1:**
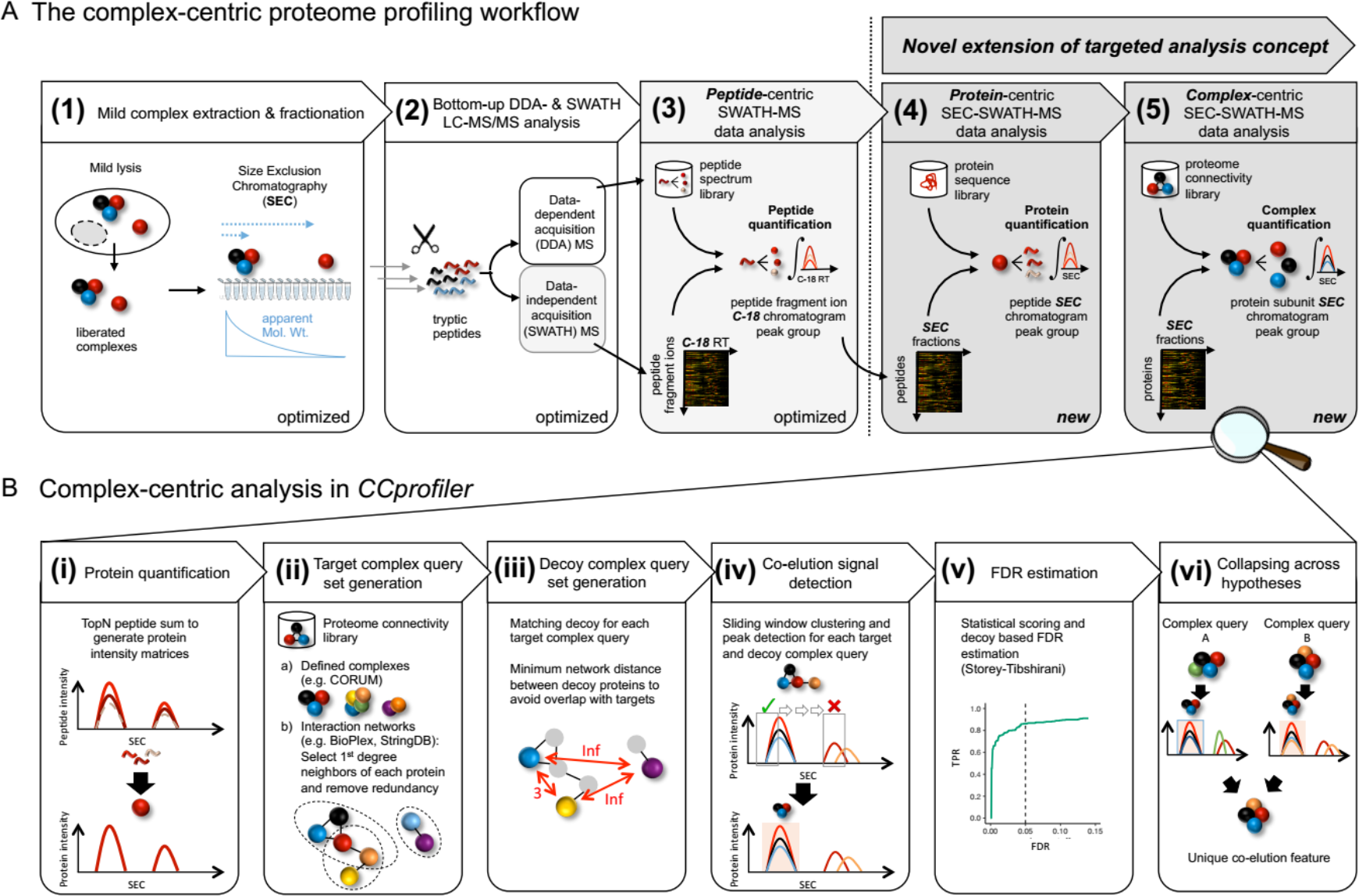
Scheme of complex-centric proteome profiling by SEC-SWATH-MS. **A** Workflow to quantify cellular complexes in five steps, extending the targeted analysis concept from peptide-centric interpretation of SWATH-MS data to the levels of protein and protein complex detection from size exclusion chromatographic fractions (also see Figure S1). **B** Specific steps of targeted, complex-centric analysis of co-fractionation data in the *CCprofiler* package.

### Benchmarking and performance assessment

We evaluated the performance of the described complex-centric analysis method, (i) by benchmarking the *CCprofiler* algorithm and error model against a manually curated reference data set, (ii) by comparing its performance with the performance of a reference method consisting of multidimensional co-fractionation of native complexes and the proteomic analysis of 1,163 fractions by data dependent mass spectrometry^12^, and (iii) by demonstrating increased sensitivity for complex detection as a result of the improved consistency of quantification of SWATH/DIA compared to data-dependent acquisition-based mass spectrometry (Figure 2).

**Figure 2:**
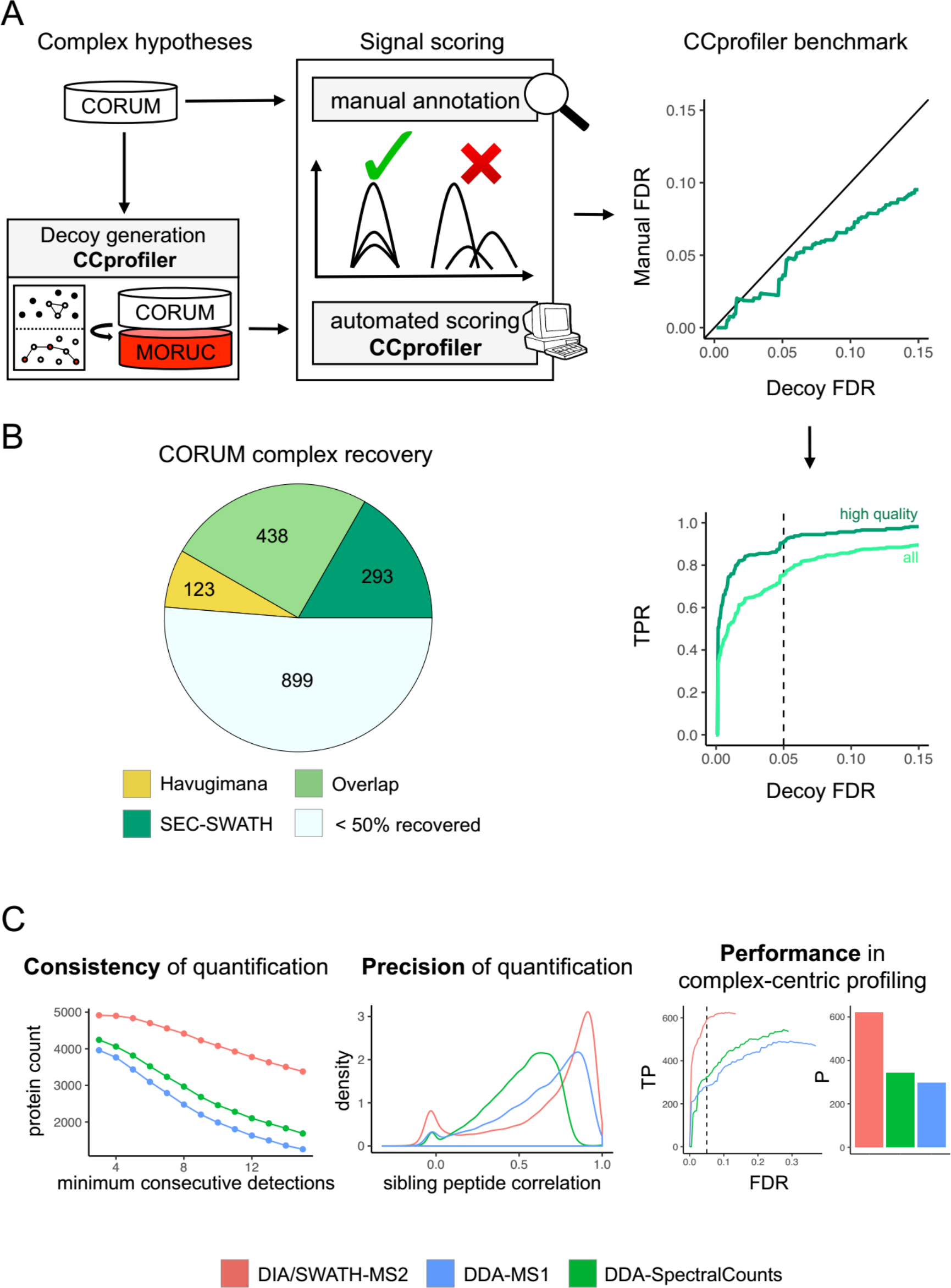
Benchmarking and performance assessment of complex-centric proteome profiling. **A** Benchmark of *CCprofiler* algorithm and error model in reference to a manually curated reference set of signals displays conservative decoy-based FDR control and high sensitivity, recalling 91 % of high quality co-elution signals (also see extended experimental procedures and Figure S2A/B). **B** Assessment of complex identification performance for the overall complex-centric profiling workflow employing CORUM, BioPlex and StringDB proteome connectivity priors, based on CORUM complex recovery in comparison to multidimensional co-fractionation and 1,176 LCMS-run-based complexome map (See Figure 4 and extended experimental procedures). **C** Comparison of SWATH-MS-based quantification to DDA-MS-based strategies (MS1 XIC and spectral counting) with regard to consistency (as judged based upon protein-level SEC chromatogram robustness towards increasing requirements on the number of consecutive detections) and precision (judged based on the correlation between sibling peptide SEC chromatograms) of quantification and overall performance in error-controlled complex-centric query of CORUM complexes in the respective protein level chromatogram sets (also see Figure S2C).

Using the data generated from the HEK293 cell line proteome, we first benchmarked the automated performance of complex-centric analysis and FDR estimation by *CCprofiler* against a manually curated reference set (Figure 2A). The manual reference set was generated by manually testing protein complexes reported in the CORUM knowledgebase^15^ for evidence of complete or partial co-elution signals among the protein level SEC chromatograms of the respective protein subunits (criteria: ≥ 2 proteins show at least one chromatographic co-elution peak, as judged by visual expert inspection). Taking the manually annotated co-elution signals as ground truth, a false discovery rate (FDR) of the complex detection in *CCprofiler* could be estimated based on the number of automatically detected complex signals that were not confirmed by the manual reference set (manual FDR, also see extended experimental procedures). This manual reference based FDR was compared to the independent FDR estimation by the target-decoy approach (decoy FDR), demonstrating that the target-decoy model provides accurate or slightly conservative error estimates of the algorithm (Figure 2A). To further evaluate the sensitivity of *CCprofiler*, we tested the recall of manually detected signals by the automated analysis. At 5% decoy-estimated FDR, the automated *CCprofiler* analyses recalled 91% of the high-confidence manual signals and 76% of all manually annotated signals (Figure 2A and Figures S2A/B).

Second, we compared the performance of the complex-centric analysis method with the performance of a reference *de novo* complex analysis method implemented by Havugimana et al. which is based on multidimensional fractionation of native complexes isolated from HEK293 and HeLa cells^12^ (Figure 2B). As a metric, we evaluated the ability of either method to recall complexes reported in the CORUM knowledgebase which consists of a total of 1753 non-redundant complexes. We considered a complex as recalled if at least 50 % of its CORUM annotated protein subunits were stated as part of a reported complex by either method (For details, see extended experimental procedures). The comparison comprised all 622 reported complexes from Havugimana et al. with unknown error rate, compared to the collapsed set of 462 complexes derived from complex-centric analysis of our HEK293 SEC-SWATH-MS dataset based on prior information from CORUM, BioPlex and StringDB, each independently filtered for 5% FDR. The results show that the complex-centric analysis method recalled 731 complexes from 81 fractions generated by single dimension SEC, compared to 561 complexes recalled from 1,163 fractions by the multidimensional reference method ^12^ (Figure 2B). The results also show a large agreement (438) between the recovered CORUM complexes. However, both datasets also uniquely recall parts of the CORUM complexes (123 complexes were uniquely confirmed by Havugimana et al. and 293 by our workflow, respectively).

Due to a lack of ground truth in terms of the set of biologically present protein complexes in each respective dataset, ultimate conclusions on the correctness of each set of reported complexes remains challenging. However, these results demonstrate that a single-fractionation mass spectrometry dataset analyzed by complex-centric proteome profiling can retrieve comparable, if not more comprehensive information on the protein complex landscape, as compared to previous multidimensional fractionation efforts including a fourteen times higher number of sample injections coupled to *de novo* complex analysis.

Third, to assess the contribution of SWATH/DIA quantification to the favorable recall results of the complex-centric proteome profiling workflow, we compared results obtained by SWATH/DIA based protein quantification with those obtained by MS1 signal integration or spectral counting when the same samples were analyzed by DDA. To generate the DDA dataset, aliquots of the peptide samples of the 81 SEC fractions analyzed by SWATH/DIA were also analyzed by data-dependent acquisition on the same TOF model 5600 mass spectrometer that was also used for SWATH/DIA acquisition. Results are shown in Figure 2C. At a respective protein-level FDR control of 1 %, SWATH/DIA quantifies 4916 proteins across the SEC fractions (≥ 2 independent proteotypic peptides), whereas the DDA data only covers 4176 proteins when analyzed by MS1 quantification based on the top2 intensity sum, and 4497 proteins when quantified by spectral counting (for details on the respective data analysis strategies see extended experimental procedures). To further assess the differences between DIA and DDA quantification, we next analyzed the three datasets with respect to the consistency of protein detection and quantification along consecutive SEC fractions (Figure 2C, left panel). The results indicate that SWATH/DIA detects and quantifies a substantially higher number of proteins in three or more consecutive fractions compared to DDA based analyses. Next, the precision of quantification as judged by global correlation among quantitative profiles of peptides originating from the same parent protein was compared between SWATH/DIA and DDA quantification, showing favorable quantification precision for SWATH/DIA (Figure 2C, middle panel). Finally, the DIA and DDA datasets were compared by their performance in detecting protein complexes (Figure 2C). At 5% controlled FDR, complex-centric analysis provides co-elution evidence for 621 versus 298 and 343 of the CORUM set of query complexes from quantitative data from SWATH-MS2, DDA-MS1 and DDA-spectral counting, respectively. Overall, these results demonstrate the favorable quantitative characteristics of SWATH/DIA data compared to DDA based analyses of data acquired on the same Triple TOF model 5600 mass spectrometer (also see Figure S2C).

The presented results demonstrate that automated complex-centric analysis by *CCprofiler* allows protein complex detection at a high sensitivity compared to manual inspection and that the system provides an accurate decoy model for FDR estimation. The data further suggests that complex-centric proteome profiling achieves competitive complex detection performance of the overall workflow with only 81 LC-MS/MS measurements compared to a significantly larger scale multidimensional fractionation experiment. Furthermore, our comparative analysis attests SWATH/DIA more consistent and precise quantification when compared to DDA-based strategies and largely increased sensitivity in targeted, complex-centric profiling under strict error rate control.

### Complex-centric analysis of the HEK293 proteome: Insights into proteome modularity

We applied the complex-centric proteome profiling method to study the modularity of the HEK293 cell line proteome. Specifically, we first used the quantitative capacity of the method to estimate the fraction of the observed proteome that was, under the extraction conditions used, part of protein complexes as opposed to being present in monomeric form. Second, we tested the ability of the method to conclusively confirm the presence of specific complexes in the sample and third, we assessed the capability of the method to quantify the distribution of specific proteins across different complexes.

#### Complex assembly state of the HEK293 proteome

To globally assess the state of assembly of the HEK293 proteome under the extraction and SEC conditions used, we quantified for each of the 4916 proteins identified in the dataset (see above and extended experimental procedures) the proportion that was detected in assembled or monomeric state, respectively. To assign a protein signal to either state, we first calibrated a molecular weight scale of proteins expected in each SEC fraction using a reference set of proteins with known molecular weight (Figure S3). We then applied this scale to all detected proteins. We assigned proteins to an assembled state if they eluted from the SEC column at an apparent molecular weight that was minimally two times higher than the molecular weight indicated by the molecular weight scale (Figure 3A). To assess the distribution of proteins across distinct molecular weight regions, indicative of different assembly states as described above, we performed a protein-centric analysis of the 58,792 peptide-level chromatograms (Figure 3A, compare Figure 1A, Step 4). Our analysis identifies 5503 elution peaks for 4065 proteins (Supplemental Table 1), with no defined elution peaks observable from the remaining 851 proteins. Of these, 2668 proteins (66 %) were observed in at least one assembled state, whereas 1397 proteins (34%) were detected only in monomeric state, based on the criteria used (Figure 3B). Of the 4065 proteins, 1103 proteins (27 %) eluted in more than one peak and up to 6 elution peaks per protein were detected (Figure 3C). Proteins that were detected in multiple assembled states were enriched in proteasome components, ribosomal proteins and chaperones (Figure 3D). We further estimated the total protein mass that was detected in assembled vs. monomeric state by integrating the total MS signals observed for proteins assigned to assembled or monomeric states. The results show that 55 % of the detected protein mass was in assembled state (Figure 3B).

**Figure 3:**
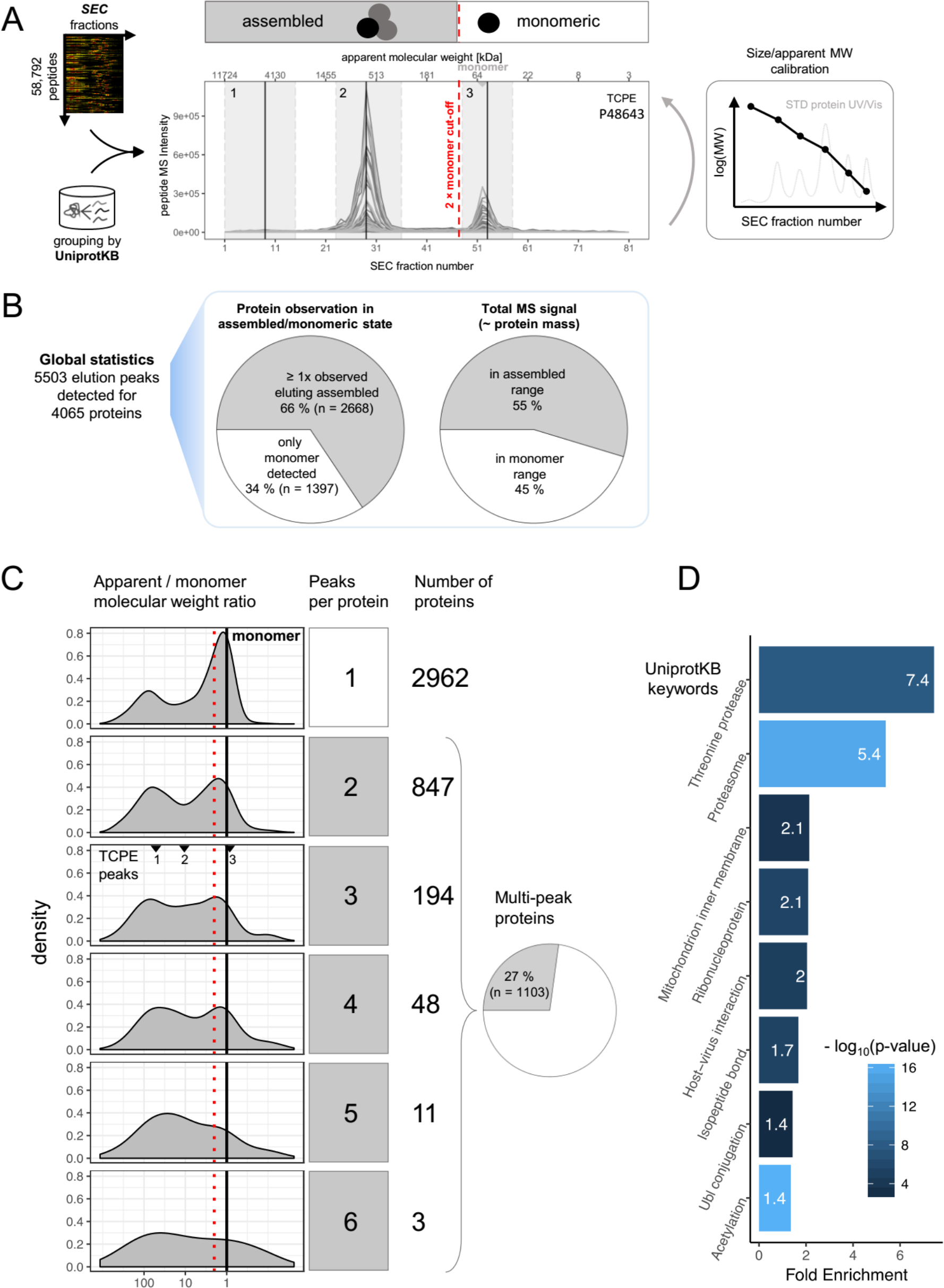
Detection of protein elution via protein-centric analysis. **A** Peptide-level SEC chromatograms are grouped by UniprotKB identifier to detect co-elution signals indicative of protein elution ranges/peaks. Based on external size calibration of the apparent analyte molecular weight per SEC fraction, signals can be attributed to likely assembled or monomeric state (also see Figure S3). For TCPE, three distinct elution signals, numbered 1-3, are detected, two in the assembled and one in the monomer elution range. **B** Global statistics of protein signal attribution to assembled or monomeric state. The majority of proteins (66 %) and protein mass (55 %) appear in assembled state in SEC-SWATH-MS. **C** Proteins are observed eluting in 1-6 distinct peaks and with a wide range of apparent vs. monomeric molecular weight ratios (distributions, left panels). The molecular weight ratios of the three peaks detected for TCPE (displayed in A) are indicated. Many of the proteins eluting in a single peak (top panel and bar) appear assembled. Proteins eluting multiple times (27 % of the proteins) do so preferentially in the assembled range, suggesting frequent participation in multiple differently sized macromolecular assemblies (lower panels and pie chart). For a list of all detected protein peaks, see Supplemental Table 1. **D** Proteins observed in multiple assembled peaks (n = 659) are enriched in components of the proteasome and other known large complex assemblies.

Overall, these results indicate that a substantial fraction of the HEK293 proteome was detected in an assembled state, both in terms of distinct protein elution peaks and protein mass (Figure 3B). The results further demonstrate the capability of the method to quantify the distribution of proteins that are part of different distinct complex assemblies (Figure 3C).

#### Complex-centric detection and quantification of complexes

As a next step we used the complex-centric workflow to confirm the presence of specific complexes in the HEK293 cell sample. The query complexes were predicted from the CORUM, BioPlex and StringDB reference databases of protein interactions, respectively, and the predictions were tested by *CCProfiler* using the 4916 protein SEC elution profiles detected in the dataset (Figure 4 and compare Figure 1A, Step 5). At a FDR of 5 % computed by the target-decoy model of *CCProfiler*, complex-centric analysis confirmed 621, 1052 and 1795 of the tested query complexes from the three respective input databases (For details, see extended experimental procedures and Supplemental Table 2). Notably, *CCprofiler* was able to confidently detect complexes consisting of the whole set of proteins predicted from the respective reference databases as well as complex signals comprising only a subset of the reference proteins, thus supporting the quantification of fully and partially assembled complexes. Up to this point in the analysis workflow, each protein complex signal detected by *CCprofiler* is directly linked to one specific protein complex query in the prior information dataset, derived from either CORUM, BioPlex or StringDB. However, some of the subunits in each complex query might overlap with other complex queries. One simple example would be that complex query A consists of subunits WXYZ and complex query B consists of subunits VXYZ. If only XYZ are detected as a co-elution group in the data, they will, until this point, be reported for both complex query A and B. In order to retrieve truly unique signals, the reported complex signals can finally be collapsed based on a strategy that considers (i) subunit composition and (ii) resolution in the chromatographic dimension. Taking the simple example from above, the signal collapsing step will merge the two features from complex query A (XYZ discovered from querying WXYZ) and B (XYZ discovered from querying VXYZ) to one unique protein signal XYZ that is independent from the original complex queries (For more details, see extended experimental procedures and Figure S4). According to this strategy, our integrated analysis across the three sets of complex queries identifies 462 unique protein-protein complex signals (Figure 4).

**Figure 4:**
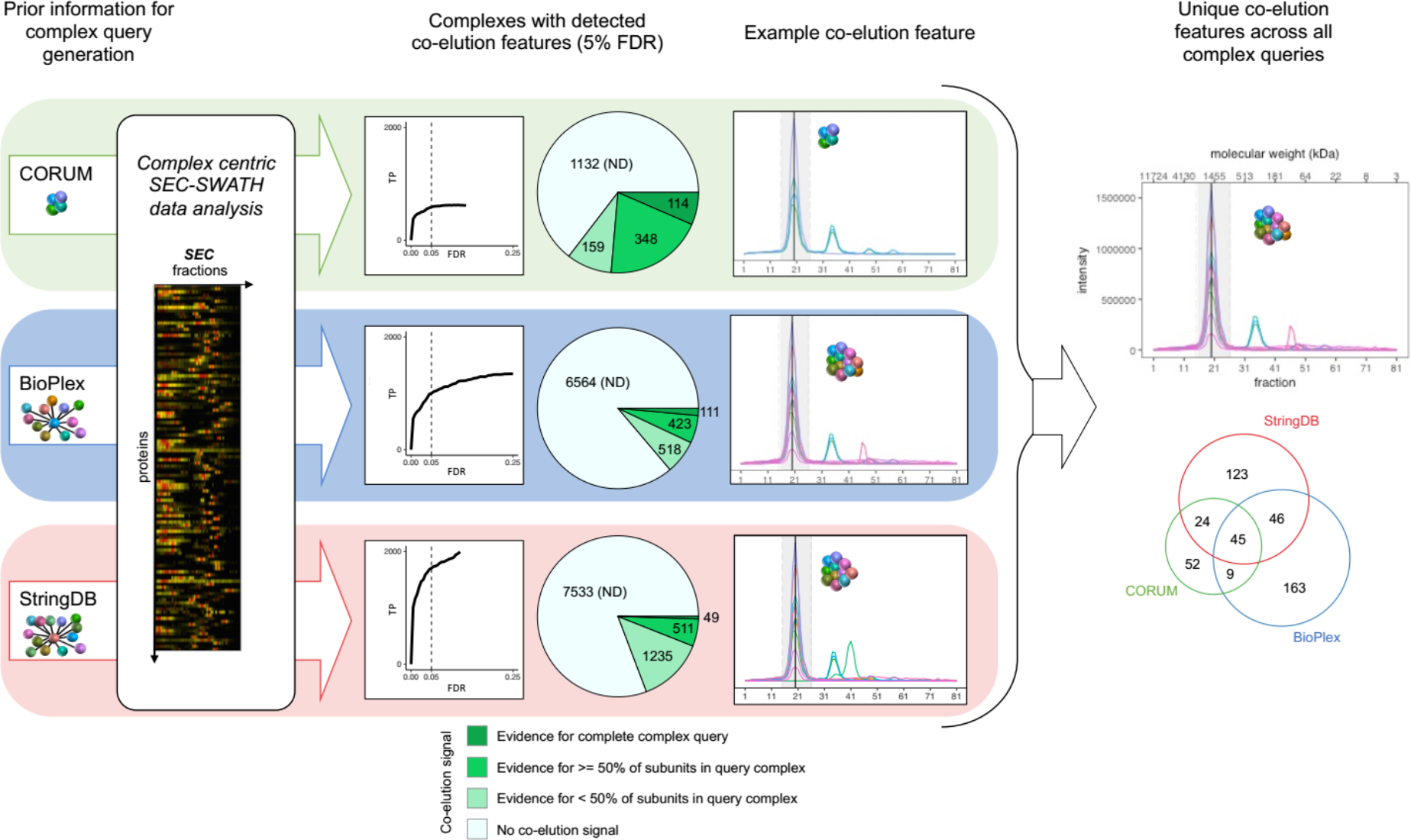
Complex-centric profiling of the HEK293 proteome by targeted query of CORUM, BioPlex and StringDB. Schematic overview of the targeted, complex-centric analysis of the protein-level cofractionation map recorded in SEC-SWATH-MS via *CCprofiler*. The three-tiered analysis is centered on complex hypotheses (i.e. groups of proteins queried for co-elution in the SEC data) obtained from CORUM or formulated from BioPlex and StringDB. At complex hypothesis FDR controlled to 5 % via the decoy-based error model, co-elution evidence is confidently detected for 621, 1052 and 1795 (representing 35.4, 13.8 and 19.2 %) of the queried CORUM-, BioPlex- and StringDB-derived hypotheses, respectively. Heterogeneity and redundancy within and across the different hypothesis sets translates to the co-elution signals retrieved, which, pieced together by collapsing on composition and SEC elution fraction, identify 462 distinct, chromatographically resolved co-elution groups representative of distinct complexes or equisized families of complexes (also see Figure S4). For a list of all detected complex signals, see Supplemental Table 2.

#### Complex-centric detection of complex variants

The results above established the capacity of complex-centric profiling to detect and quantify subunit distribution across complexes that are resolved by SEC and contain common proteins. We therefore tested whether this capacity allowed us to detect novel protein modules of potential functional significance. Among the 621 complex models that were confirmed by CCProfiler following predictions from the CORUM database, 286 (46%) provided evidence of proteins common to two (152) and up to 5 or more (27) distinct, chromatographically separated complex instances (Figure 5A). For example, the protein subunit fractionation profiles of the octameric COP9 signalosome complex, a central regulator of E3 ligase activity and turnover, delineate both the CSN holo-complex consisting of all eight subunits and a sub-complex consisting of subunits CSN1, CSN3 and CSN8 (Figure 5A&B). The critical role of CSN proteins in regulating the ubiquitin-proteasome system and cellular homeostasis has sparked great interest in the analysis in modules with variable subunit composition, and in mechanisms that regulate their activity ^20^. CSN proteins have also been linked to cancerogenesis^21–23^. Both CSN assemblies detected in the HEK293 dataset elute with apparent molecular weights in accordance with a 1:1 stoichiometry. Further, the proteins CSN1/3/8 of the lower molecular weight complex form a connected sub-module within the CSN holo-complex structure^24^ (Figure 5C). The occurrence of the distinct CSN1/3/8 complex detected in this study is consistent with protein chromatographic data generated by co-fractionation-MS/MS in two other laboratories. Wan et al. fractionated mild lysates of HEK293 cells by heparin ion exchange chromatography^25^ followed by MS analysis. Also in this separation dimension that is orthogonal to SEC, the quantitative MS profiles show distinct co-fractionation of CSN1, CSN3 and CSN8 (Figure 5D, upper two panels). Larance et al. fractionated mild lysates of U2OS cells by SEC and the quantitative profiles of the CSN subunits also display distinct co-elution of CSN1, CSN3 and CSN8 at reduced molecular weight (Figure 5D, lower panel). While the data of both research groups generally support the model of CSN1/3/8 as a distinct cellular assembly, neither of them reported it as distinct from CSN holo-complex, likely owed to limited resolution of the experimental data and the pairwise-interaction-focused analysis workflows employed. Our findings suggest a potential functional role for the CSN sub-complex CSN1/3/8. We therefore tested whether CSN1, CSN3 and CSN8 could stably assemble independent of the remaining CSN components. We co-expressed human CSN1, CSN3 and CSN8 in insect cells, whereby CSN8 was added with an N-terminal Strep(II)-tag and CSN1 & CSN3 were expressed with and N-terminal His6-tags to facilitate reciprocal purification of the complex. The thus purified samples were analyzed by SDS-PAGE and resulting banding patterns confirmed the formation of a stable trimer CSN1/3/8 in the absence of the other CSN subunits that constitute the holo-complex (Figure 5D). Together, these results support the finding from the complex-centric identification of the CSN1/3/8 complex as a distinct substructure of the human COP9 Signalosome.

**Figure 5:**
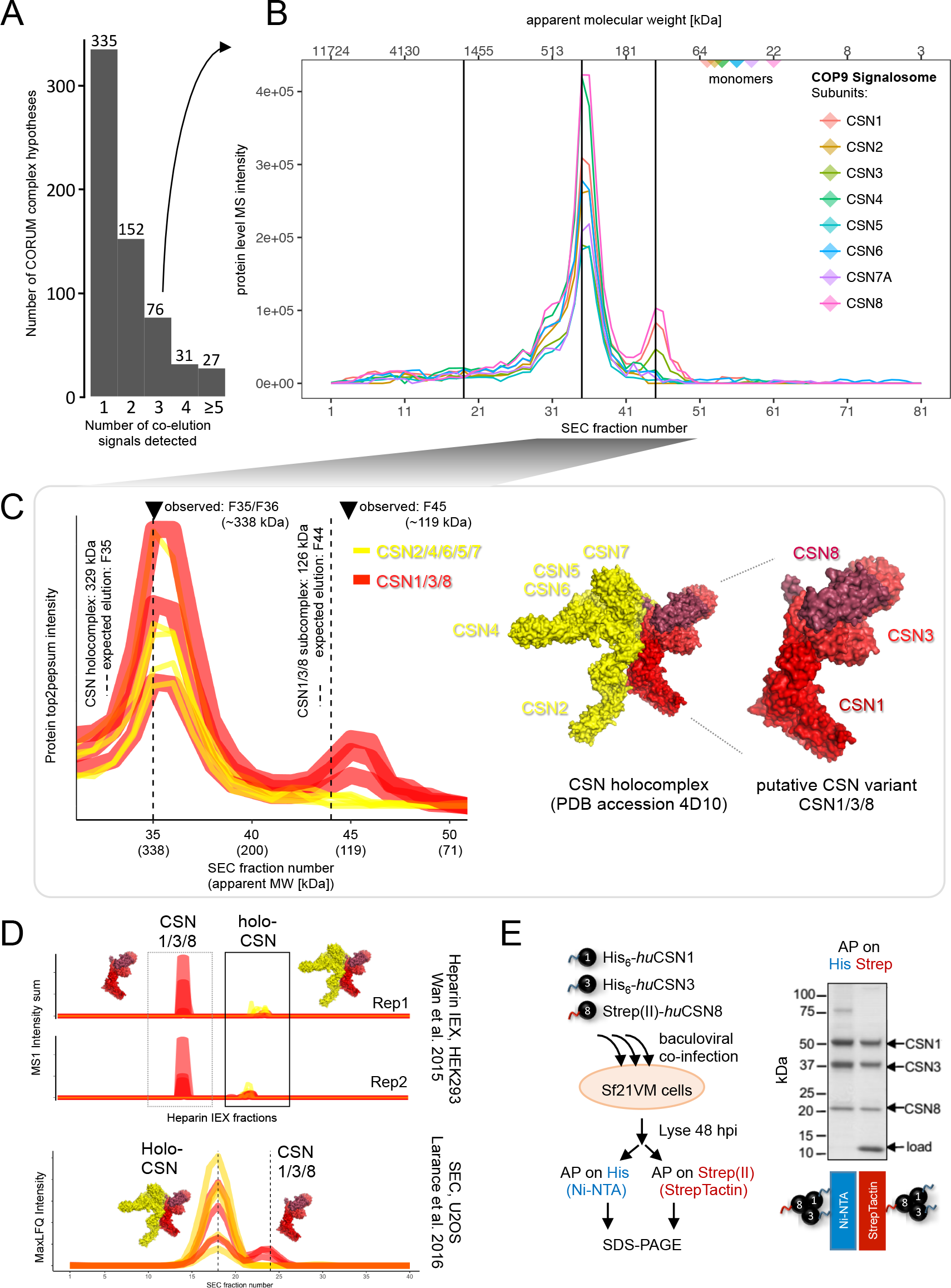
Complex-centric detection of COP9 Signalosome variant CSN1/3/8. **A** For nearly half the CORUM complex hypotheses queried, two or more distinct subunit co-elution signals were detected (See extended experimental procedures). **B** Among the four distinct co-elution signals detected from the eight canonical CSN subunits’ chromatograms (here with CSN7A, not CSN7B) are two distinct signals delineating **C** distinct co-elution of holo-CSN (observed at the expected fraction 35) and Mini-CSN CSN1/3/8 (observed eluting offset only one fraction late, F45, of the expected fraction, F44). Expected fractions are estimated from the cumulative sum of one copy per component and external size calibration. Coloring adapted to highlight subversion components and their partitioning across holo- and sub-complex. CSN1/3/8 interact and form a submodule within the CSN holo-complex structure (PDB accession 4D10). **D** CSN1/3/8 display distinct fractionation patterns in co-fractionation experiments performed in other laboratories, specifically in orthogonal ion exchange fractionation of HEK293 lysates (Wan et al. 2015, upper panels) and size exclusion chromatographic fractionation of U2OS lysates (Larance et al. 2016, lower panel), in line with the CSN1/3/8 as distinct entity.

As a further example for the discovery of sub-complexes of a large holo-complex, complex-centric proteome profiling detected six variant signals from the subunit chromatograms of the 26S proteasome (Figure 6A). Two of the six variants represent known complexes, (i) the full 26S assembly and (ii) the 20S core particle (Figure 6A). The remaining four co-elution signals point towards complex variants of lower apparent molecular weight compared to the 26S and 20S particles (apex fractions 39, 40, 42 and 46, ^~^ 107-222 kDa) that consist predominantly of α and β subunits of the 20S core particle. These reported complex variants point towards (iii) a β subunit assembly of β2, β3 and β7 at fraction 39, (iv) a distinct assembly of α subunits α2 and α6 at fraction 40, (v) an assembly intermediate of the seven α subunits α1, α2, α3, α4, α5, α6 and α7 at fraction 42, and (vi) a β6 and proteasome regulatory subunit 8 assembly at fraction 46. To evaluate whether the observed signals represent products of disassembly or complex biogenesis intermediates, we manually extended the automated analysis of CCProfiler by additionally aligning the quantitative protein traces of the chaperones known to be involved in 20S maturation with the respective complex subunits^26^(Figure 6B). Strikingly, the distinctive co-elution of the early-stage specific chaperone PSMG3/PSMG4 dimer, constitutive chaperone PSMG1/PSMG2 dimer and the late-stage specific proteasome maturation factor POMP allowed us to classify the detected complex variants as early and late stage intermediates of 20S biogenesis (Figure 6B). Notably, a systematic manual analysis of the quantitative distribution of the proteasome and chaperone subunits across the detected complex variants suggests the α1/α3/α4/α5/α7 and α1-7/β2/β3/β6/β7 complexes, respectively, as the predominant early and late assembly intermediates on the path to 20S assembly, as assigned by defined co-elution and inferred interaction with the chaperones specifically involved in early (PSMG3/PSMG4 dimer) stages or late stages (POMP) of 20S proteasome biogenesis (Figure 6C). Although the automated workflow could not fully resolve and explain the data, it successfully pointed towards a distinct assembly of the alpha subunits (signal v) from the beta subunits (signal iii), as well as the differential behavior of α2 and α6 compared to the other α subunits (signal iv). No underlying biology could be determined for signal vi.

**Figure 6:**
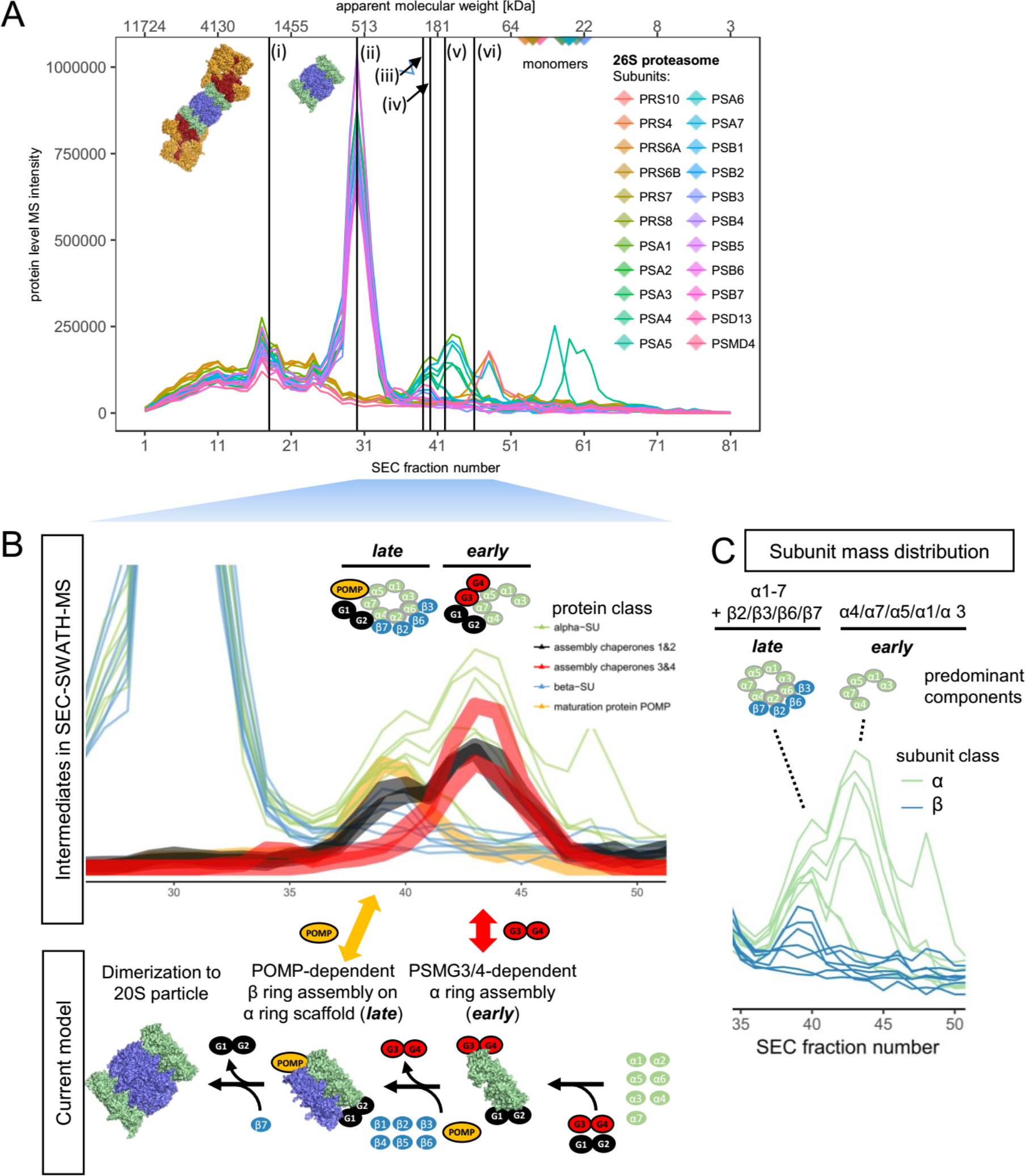
Complex-centric detection of 20S proteasome assembly intermediates. **A** Protein-level SEC chromatograms of the 22 canonical 26S proteasome subunits. Vertical black lines indicate the apexes of six distinct co-elution signals detected in complex-centric scoring; two of which represent well-known co-occurring variants, the full 26S (i) and the 20S (ii) particle devoid the 19S lid and ATPase (Indicated by structural models, PDB accession 5GJR) and four of which, composed of predominantly 20S a and P subunits, appear at reduced size (222 – 107 kDa, fractions 39, 40, 42 and 46). **B** Zoom in to chromatograms of 20S components in full and reduced MW range and in the context of chaperones known to be involved in assembly according to the current model of 20S biogenesis (lower panel, assembled after Saeki & Tanaka, 2012^38^ and PDB accession 5GJR), colored by protein class. Reduced MW species are classified into early and late assembly intermediates (as opposed to artifacts of disassembly) by defined co-elution of early assembly chaperone PSMG3/PSMG4 dimer, late assembly chaperone proteasome maturation protein POMP and constitutive chaperone PSMG1/2 dimer. **C** Subunit mass distribution across early and late assembly intermediate elution ranges suggests predominant components of the intermediary species accumulating in HEK293 cells.

Together, these findings demonstrate the capacity of complex-centric profiling to derive models of distinct variants of the queried complexes. These models can be reinforced by extending automated analyses by the alignment of additional proteins’ quantitative profiles followed by manual inspection.

#### SECexplorer – An interactive platform for complex-centric exploration of the HEK293 proteome analyzed by SEC-SWATH-MS

To support customized, expert-driven and in-depth analyses of protein co-fractionation profiles recorded by SEC-SWATH-MS of the HEK293 cell line, we set up the web platform *SECexplorer*. *SECexplorer* enables visualization and interactive browsing of protein fractionation profiles of user-defined sets of proteins. Users can perform multiple tasks, including (i) testing of novel predicted models on complex formation between candidate proteins or (ii) interrogating the profile sets of known modules for evidence pointing towards new variants or (iii) manual refinement and extension of results obtained from automated complex-centric profiling, for example by extending the set of automatically detected complex components with additional proteins e.g. derived from the literature or from interaction network context. Analyses are assisted by the *CCprofiler* algorithm suggesting distinct co-elution signals and calculating their expected to apparent molecular weight mismatch, among other metrics, in order to speed up data interpretation by expert users. *SECexplorer* can be accessed at *https://sec-explorer.ethz.ch/* (Figure 7A). As an example for the use of *SECexplorer*, we followed up on the peak shoulder at elevated molecular weight observed in the CSN holo-complex co-elution signal (Compare Figure 5B). Overlaying the elution profiles of known components of a E3-CRL substrate of the COP9 Signalosome^27^ revealed defined co-elution in the peak shoulder range, supporting the detection of a likely E3-CRL-bound subpopulation of CSN holo-complexes (Figure 7B). To derive a quantitative signal in the situation of only partial chromatographic resolution we employed a Gaussian deconvolution mixture model, suggesting a substrate-bound fraction of CSN holo-complex of 25 ± 3 % across the 8 subunits (Figures 7C and S5).

**Figure 7:**
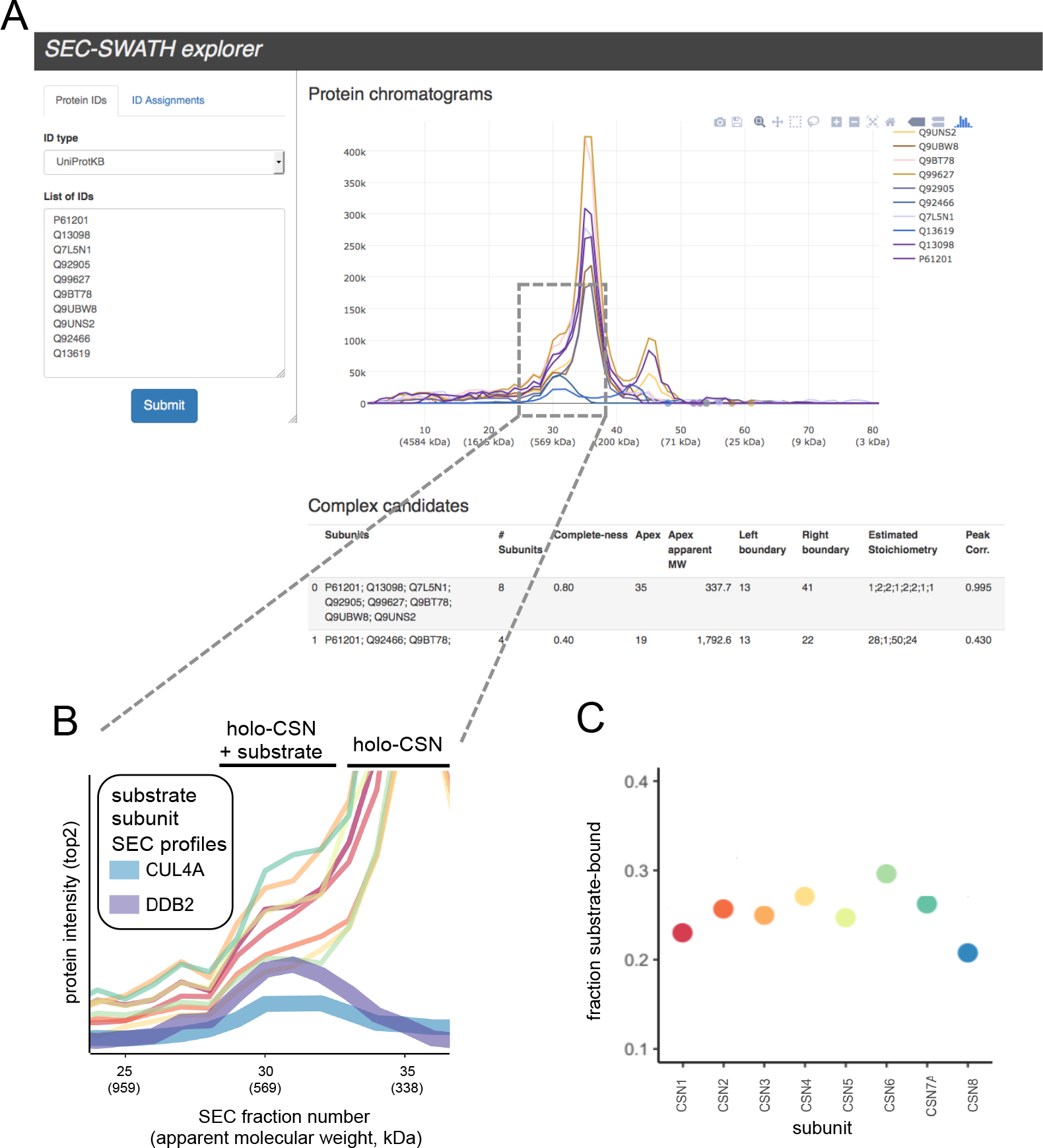
SECexplorer tool for customized interrogation of SEC-SWATH elution profiles. **A** SECexplorer web-interface for querying custom protein sets for co-elution behavior in the SEC-SWATH-MS data, viewing chromatograms for interpretation and with algorithmic assistance. **B** Zoom in to high MW peak shoulder of holo-COP9 Signalosome (compare Figure 5), where defined coelution signals of CSN substrate components CUL4A and DDB2 suggest the partial resolution of substrate-bound and free pools of CSN holo-complex. **C** Estimation of the fraction of holo-CSN in the likely substrate-bound pool versus the free pool, with eight measurements along the eight subunits and based on Gaussian deconvolution of two signals underlying the observed peak and shoulder (also see Figure S5).

## Discussion

In this paper we describe complex-centric proteome profiling, an integrated experimental and computational approach to detect and quantify protein complexes isolated from their natural source, to generate new insights into the modular organization of proteomes.

The need to systematically analyze the organization of the proteome arises from the notion of a modular biology proposed by Hartwell et al.^1^. It essentially states that biochemical functions are for the most part catalyzed and controlled by functional modules, most frequently protein complexes, and that (genomic) perturbation of complexes results in perturbed biochemical functions and potentially in disease phenotypes. The notion of a modular biology thus extends the pioneering work of Pauling et al. on defining sickle cell anemia as a molecular disease^28^ to the proteome level. Protein complexes and protein-protein interactions have been studied extensively by a wide range of techniques and have led to compendia of complexes^15,16,29^ and maps of protein interaction networks^16,30,31^. These compendia have in common that they describe generic, usually static instances of complexes and interactions^32–34^. To distinguish between different biochemical states of a cell, it is also essential to determine qualitative and quantitative differences in functional modules in different samples. To date this has been attempted by two broad approaches. The first is based on microscopic methods including FRET^35^ which provide outstanding resolution and precision of steric proximity but are labor intensive and focused on one to a few interactions at a time. The second is based on a mass spectrometric approach referred to as correlation profiling^36^ in which samples of native modules are separated into a set of fractions and the protein contents of each fraction is determined by quantitative mass spectrometry. The association of a protein to a specific module is then asserted by the consistency of the quantitative pattern of the protein in question with other proteins of the same module^37^. Initially used to define the composition of the specific modules such as the large RNA polymerase II preinitiation complex^37^ and the human centrosome ^38^, correlation profiling has also been employed to broadly assign protein localization to different subcellular compartments^39–41^and the scope has been extended towards systematically interrogating protein-protein complexes by correlating protein patterns in fractions obtained from different biochemical fractionation methods^9,11,42^. Such studies have used different native complex separation methods including SEC, IEX, density gradient centrifugation and blue native gels^9,11,42^. The scientific scope has extended to the analysis of cells of different species, culminating in the description of hundreds of complexes in a single, albeit massive experiment^25^. Correlation profiling, therefore has the potential to determine the quantity and composition of hundreds of protein modules in a single operation.

In the present paper we describe a conceptual and technical advance in the field of correlation profiling. As a conceptual advance we introduce the principle of complex centric analysis. It is inspired by the peptide-centric analysis concept employed for the specific and sensitive detection of peptides from proteomic samples in targeted proteomic approaches, such as SWATH/DIA^6^, and extends the use of prior information for the analysis of proteins to the level of protein complexes. Similar to peptide-centric analysis of SWATH/DIA data, high selectivity and sensitivity is achieved by focusing the analysis on analytes conceivably expected in the sample when querying the protein co-fractionation data for candidate protein complexes that are inferred from reference protein interaction maps. Thereby, prior information significantly constrains complex inference from cofractionation profiling data and thus adds specificity and the possibility to develop a target-decoy model to assess the reliability of the obtained results. Furthermore, complex identifications are directly linked to quantitative chromatographic signals, a central feature of targeted proteomics approaches. As technical advances we demonstrate the benefits of SWATH/DIA for the analysis of the sequential SEC fractions, introduce a freely accessible computational framework *CCprofiler* and provide a tool facilitating the exploitation of complex-centric data, *SECexplorer* (Figure 7).

In combination, these technical and conceptual developments provide the following advances to the field of correlation profiling. First, the preferable quantitative performance of SWATH/DIA provides more complete and consistent sampling of the eluting proteome, resulting in fewer gaps and noise in the recorded profiles. This results in deeper insights into modular proteome organization, including the detectability of low abundance complex intermediates. Second, the use of prior information reduces false positive assignments of complex co-membership due to coincidental co-elution of proteins that do in reality not interact. Third, the *CCprofiler* pipeline introduces the first statistical target-decoy model to tightly control error rates in the inference of complexes from co-fractionation profiling experiments and represents a comprehensive, open-source platform to support complex-centric profiling of proteomes, irrespective of the fractionation method used. Fourth, the efficiency of information retrieval and thus overall method throughput is drastically increased when compared to current co-fractionation based complex analyses, generating comprehensive and accurate assessments of proteome arrangement from an order of magnitude less LC-MS experiments than necessitated earlier. Together, these advances transform the SWATH/DIA-based complex-centric proteome profiling into a robust, generally applicable technique supported by a freely accessible computational framework.

We applied complex-centric profiling to a native protein extract from exponentially growing Hek293 cells. Collectively, the results demonstrate the superior performance of the technique compared to the state of the art and provide new biological insights, as follows. The analysis establishes estimates for the overall assembly state of a human proteome - x55 % of inferred protein mass and two thirds (66 %) of the observed protein species appear engaged in higher order assemblies; a lower-boundary estimate given inevitable losses of associations in the experimental procedure. Besides detecting cumulatively 462 cellular complexes upon targeted analysis, the method in many instances resolves distinct variants of the expected complexes, such as sub-complexes that elute independently from the chromatographic column. While sub-complex signals may originate from artefactual disruption of cellular complexes, we demonstrate in two cases how orthogonal pieces of evidence can build confidence in the biological relevance of substructures assigned from defined subunit co-elution. First, we identified a new complex CSN1/3/8 as a sub-complex of the COP Signalosome (CSN) holocomplex that elicits crucial regulatory functions towards E3 ligase complexes and the ubiquitin proteasome system^43^. It is tempting to speculate that a putative function of the CSN1/3/8 subcomplex could be the negative regulation of CSN holocomplex activity, due to the fact that the subcomplex incorporates the subunit CSN1 which is involved in substrate recognition^27^, but does not contain the catalytically active CSN5 subunit. CSN5 embodies the de-neddylation activity to the CSN holocomplex^27^. CSN1/3/8 may potentially sequester neddylated E3 CRLs from CSN-mediated de-neddylation and thus affect their lifetimes and overall activity profiles. In a second example, complex-centric analysis in combination with manual refinement identified early and late assembly intermediates on the path towards the 20S proteasome particle. Strikingly, the early and late intermediary complexes assigned (early: α1/α3/α4/α5/α7, late: α1-7/β2/β3/β6/β7) collide with current models of the temporal order of subunit assembly^44,45^ (for a graphical summary, see Figure 6B, lower panel). Current models entail early α-ring intermediates lacking subunits α3 and α4^46^. In contrast, our model suggests assembly of pre-α-ring intermediates composed of subunits α4, α7 α5 α1 and α3 (forming a connected substructure of the α-ring in this order^47^) that lacks subunits α2 and α6. These join thereafter to complete the α-ring, under involvement of the chaperone POMP/hUmp1. Current models further suggest that ordered β-ring assembly scaffolded by α-rings in the sequence of β2, β3, β4, β5, β6, β1 and lastly β7^44,45^ help overcome a POMP-dependent checkpoint for dimerization into the mature 20S particle^48^. The detection of late assembly intermediate α1-7/β2/β3/β6/β7 in our data suggests an alternate sequence of assembly with early incorporation of subunit β7 and dimerization after the recruitment of subunits β1, β4 and β5.

These insights into complex biogenesis could prove valuable, for example, in the design of future therapeutic strategies aiming to counteract elevated proteasome expression and activity that has been associated with cancer pathobiology^49^. This is exemplified by current attempts to target proteasomal activity via the chaperone POMP^50^ (Figure 6C). We expect that the data generated by complex-centric proteome profiling will lead to the discovery of other instances of characteristic protein complexes and sub-complexes and thus trigger research into their functional roles.

Despite the advances and benefits of complex-centric proteome profiling by SEC-SWATH-MS, the method has a number of limitations. i) The balance of stability of complexes and extractability in native form. Inevitably, associations are lost in the experimental procedure, most notably upon dilution imposed during lysis and subsequent size exclusion chromatography, reducing protein concentration by ca. five orders of magnitude from the cellular environment (ca. 300 mg/ml^51^) to the conditions on the SEC column (ca. 0.06 mg/ml). Consequently, complex detectability is limited by thermodynamic stability and despite best efforts towards minimizing complex disintegration (fast processing in the cold and analyte adsorption-free chromatography), thermodynamically labile interactions, particularly those with fast off-rates, are likely inaccessible by correlation profiling methods, including complex-centric proteome profiling. While first studies have evaluated chemical crosslinking as means to stabilize cellular modules for chromatographic analyses^52^, it remains an open challenge to identify uniformly beneficial crosslinking reagents and reaction conditions that yield optimal balance between stabilization of biologically relevant structures and artefactual cross linking across the full range of protein expression in the cell^53^.

Furthermore, complex-centric proteome profiling is limited to the scope of the prior knowledge on protein association employed. However, continued efforts to map cellular protein association space^54^ and computational integration of multiple lines of experimental evidence^29^ will continually improve the quality and completeness of the prior knowledge useable as input to targeted, complex-centric analyses. Extended reference protein interaction maps will support near-complete mapping of the complexes detectable in co-fractionation experimental data in the near future, supported by scalability of the target-decoy statistical model. That being said, the statistical model itself is limited to the assignment of an FDR on the evidence of detection of defined complexes in the complex query set. Future improvements could potentially support a robust statistical model covering also postprocessing steps, such as collapsing of detected features across multiple complex query sets to unique co-elution signals.

SEC-SWATH-MS accelerates the mapping of cellular complexes. Whereas the method yields a similar coverage of complexes compared to state-of-the art at over fourteen times less LC-MS injections, it still required 81 fractions to be analyzed at 2 h gradient time per fraction, culminating in 162 h of net MS acquisition time. This fact limits the scope for cohort studies. However, this issue may well be alleviated soon, given anticipated improvements SWATH/DIA sample throughput with minimal loss of protein coverage that seem achievable because in SWATH/DIA acquisition the number of analytes quantified does much less strongly depended on gradient length than is the case for DDA acquisition. As a consequence of the high quantitative quality of the data, the increased efficiency of the method and the error model this study lays the foundation to conclusively quantify changes of proteome organization as a function of cell state. Ultimately, extensions of our workflow will support the detection of subtle re-arrangements within proteomes that occur in response to perturbation or along central biological processes. Such insights will help foster our understanding of the importance of higher order organization of the parts to convey plasticity and regulation to cellular systems.

## Author contributions

Conceptualization, R.A., M.G., B.C.C. and M.H.; Methodology, M.H., I.B., G.R.; Software, I.B., M.H., G.R., R.H., M.F. and A.B.E.; Validation, M.H. and I.B.; Formal Analysis, M.H. and I.B.; Investigation, M.H.; Resources, M.G. and R.A.; Data Curation, I.B. and M.H.; Writing – Original Draft, M.H. and R.A.; Writing – Review & Editing, I.B., M.F., G.R., R.H., A.B.E., B.C.C., M.G. and R.A.; Completion Submitted Manuscript, M.H., I.B. and R.A.; Visualization, M.H., I.B. and R.H.; Supervision, B.C.C., M.G. and R.A.; Project Administration, M.H.; Funding Acquisition, R.A.

## Acknowledgments

We thank all Aebersold and Gstaiger lab members for helpful discussions, and with special emphasis Betty Friedrich, Audrey van Drogen, Peter Blattmann, Hannes L. Roest, Ludovic Gillet and Yansheng Liu. We further thank Prof. Nicolas Thomä, Dr. Lingaraju Manjappa of the Friedrich Miescher Institute for Biomedical Research as well as Dr. Martin Renatus and Arnaud Decock of the Novartis Institutes for Biomedical Research for materials and guidance in COP9 signalosome subunit co-expression and co-purification experiments. We would also like to thank the Scientific IT Support (ID SIS) of ETH Zurich for support and maintenance of the lab-internal computing infrastructure (iPortal) and specifically Uwe Schmitt for his help with the SECexplorer setup. The project was supported by the SystemsX.ch projects PhosphoNetX PPM and project TbX to R.A., and the European Research Council (ERC-20140AdG 670821 to R.A.). M.H. was supported by a grant from Institut Mérieux. B.C.C. was supported by a Swiss National Science Foundation Ambizione grant (PZ00P3_161435). A.B.E. was supported by the National Institutes of Health project Omics4TB Disease Progression (U19 AI106761). I.B. was supported by the Swiss National Science Foundation (grant no. 31003A_166435).

## Conflict of Interests

The authors declare that they have no conflict of interest.

## Experimental Procedures

### Preparation of native HEK293 proteome and fractions for MS analysis

HEK293 cells were obtained from ATCC and cultured in DMEM containing 10 % FCS and 50 μg/mL penicillin/streptomycin to 80 % confluency. Ca. 7e7 cells were mildly lysed by freeze-thawing into 0. 5 % NP-40 detergent- and protease and phosphatase inhibitor containing buffer, essentially as described^55^, albeit without the addition of Avidin. Lysates were cleared by 15 minutes of ultracentrifugation (100,000×g, 4 °C) and buffer was exchanged to SEC buffer (50 mM HEPES pH 7.5, 150 mM NaCl) over 30 kDa molecular weight cut-off membrane at a ratio of 1:50 and concentrated to 25-30 mg/ml (as judged by OD280). After 5 min of centrifugation at 16.9 ×g, 4 °C, the supernatant was directly subjected to fractionation on a Yarra-SEC-4000 column (300×7.8 mm, pore size 500 Å, particle size 3 μm, Phenomenex, CA, USA). Per SEC run, 1 mg native proteome (by OD280) was injected and fractionated at 500 μl/min flow rate at 4 °C, collecting fractions at 0.19 min per fraction from 10 to 28 minutes post-injection, fractions 3-83 of which were considered relevant proteome elution range and considered for further analysis with fractionation index 1-81. The fractions collected from two consecutive SEC fractionations of the same extract (2×1 mg) were pooled for subsequent bottom-up proteomic analysis. Apparent molecular weight per fraction was log-linearly calibrated based on column performance check protein mix analyzed prior and after each experimental replicate (AL0-3042, Phenomenex, CA, USA). An aliquot of the unfractionated mild proteome extract was included in peptide sample preparation and LC-MS analysis. Proteins were proteolyzed to peptide level by trypsin digestion (Promega V5111) in the presence of 1 % sodium deoxycholate (Sigma-Aldrich D6750), reduced, alkylated and de-salted on C18 reversed phase (96-Well MACROSpin Plate, The Nest Group, MA, USA) and each sample was supplemented with equal amounts of internal retention time calibration peptides (iRT kit, Biognosys, CH).

### Baculoviral co-expression and co-purification

Sf21VM Cells were maintained in ExCell420 Medium in Erlenmeyer culture flasks shaking at 27.5 °C. Human COP9 Signalosome subunits bearing N-terminal Strep(II) or His6 tags were co-expressed by co-infection of Sf21VM cells with three baculoviral vectors obtained from Lingaraju et al.^24^. After 48 h, cells were mildly lysed and COP9 signalosome subunits and complexes differentially affinity-purified on StrepTactin and Ni-NTA-coated magnetic beads (Qiagen) followed by bead boiling in SDS loading buffer and subunit detection via SDS PAGE and InstantBlue staining (Expedeon). Subunits were identified by size and in reference to individual expression and in-gel detection.

### MS analysis

LC-MS analysis of peptide samples was performed in both DDA and SWATH/DIA acquisition mode on an AB Sciex TripleTOF 5600+ instrument (AB Sciex, MA, USA), side-by-side per sample, sliding from early to late-eluting fractions. On-line reversed phase chromatography fractionated peptide samples delivering at 300 nL/min flow a 120-min gradient from 2–35% buffer B (0.1% formic acid, 90% acetonitrile) in buffer A (0.1 % formic acid, 2% acetonitrile) on a self-packed picoFrit emitter packed with 20 cm column bed of 3 μm 200-Å Magic C18 AQ stationary phase, essentially as described^6,55^. In data-dependent acquisition (DDA), MS1 survey spectra were acquired for the range of 360-1,460 m/z with a 500 ms fill time cap. The top 20 most intense precursors of charge state 2–5 were selected for CID fragmentation and MS2 spectra were collected for the range of 50–2,000 m/z, with 100 ms fill time cap and dynamic exclusion of precursor ions from reselection for 15 s, essentially as described^55^.

Data-independent acquisition (SWATH/DIA) mass spectrometry was performed using an updated scheme of 64 variably sized precursor co-isolation windows optimized for human cell lysate MS signal density (SWATH^®^ 2.0, essentially as described^56^). SWATH cycles (64 × 50 ms accumulation time) were interspersed by MS1 survey scans for the range of 360–1,460 m/z with a 250 ms fill time cap, resulting in an overall period cycle time of 3498 milliseconds. The MS2 mass range was set to 200 – 2000 m/z.

### Data processing

DDA-MS data were processed using the MaxQuant software package (version 1.5.3.17) with the human canonical SwissProt reference database (build Aug-2014), standard parameters and variable methionine oxidation and N-terminal acetylation enabled. Match between runs was enabled to facilitate ID transfer and more consistent MS1 quantification (from and to) between adjacent fractions. Raw peptide MS1 intensities of individual peptide precursor signals were further considered.

The SWATH/DIA-MS data were analyzed via targeted, peptide-centric analysis, querying > 200,000 precursors based on the combined human assay library (CAL) in the SWATH fragment ion chromatograms, using a modified OpenSWATH^7^, PyProphet^57,58^ and TRIC^18^ workflow and the iPortal framework^59^ to recover precursors at an experiment-wide assay-level (TRIC target) FDR of 5 %. Precursor-level results were summed per peptide and further filtered on chromatography-informed scores, controlling the false discovery rate on protein level to below 1 %, employing the simple target-decoy method as implemented in *CCprofiler*. The protein-level fraction of false targets was estimated using R/SWATH2stats^60^. Specific filtering rules were, first, eliminating all values part ofconsecutive identification stretches below length three and, second, exclusion of peptides based on their quantitative fractionation pattern’s average dissimilarity to those of sibling peptides (originating from the same parent protein, discarding peptides with average sibling peptide correlation coefficient (spc) below 0.316 to achieve an estimated FDR of < 1 % among the remaining 4958 proteins (4916 of which quantifiable with ≥2 proteotypic peptides for downstream analyses).

Protein-centric analysis was performed within the *CCprofiler* framework, detecting protein elution signals among peptide SEC chromatogram sets grouped by parent SwissProt protein identifier (For details, see extended experimental procedures).

Complex centric analysis was performed within the *CCprofiler* framework, detecting complex elution signals among protein SEC chromatogram sets, defined by reference databases CORUM, BioPlex and StringDB (Compare Figure 1 and Figure 3). Protein level chromatograms were generated by summing the two highest intensity proteotypic peptide chromatograms. Complex queries as well as decoy complex queries were formulated from the reference databases by splitting variants and re-merging redundant human CORUM complexes (N = 1753 query groups) and by partitioning networks into query groups of one seed protein including all direct interactors at the confidence thresholds (BioPlex interaction probability ≥ 0.75 and StringDB interaction score ≥ 900). Decoy complex queries of matching number and size were formulated in reference to the input connectivity information, ensuring that decoy complex queries do not entail interacting subsets. Co-elution signals were then detected using the signal processing algorithm of *CCprofiler* in grid search mode over a range of processing parameter sets. Parameters and results were then selected to ensure precision of ≥ 95 % among the protein peak group signals, followed by summarizing signals to unique groups at defined chromatographic retention times, chromatographically resolved complexes and complex variants.

Generation of true positive reference sets of co-elution signals To guide algorithm development and to evaluate extendibility of the target-decoy approach to the level of protein complexes, we manually curated protein subunit co-elution signals among the 1753 protein chromatogram sets defined by the CORUM complexes.

A more comprehensive and detailed protocol for all data processing and analysis steps is provided in the extended experimental procedures. The mass spectrometry proteomics data have been deposited to the ProteomeXchange Consortium (http://proteomecentral.proteomexchange.org) via the PRIDE partner repository^61^ with the dataset identifier PXD007038. The *CCprofiler* package is freely available on GitHub (https://github.com/CCprofiler/CCprofiler/) including a vignette describing the main functionalities and usage of the software (Supplemental item 1).

